# SPIFEE -- A pipeline for analyzing traces of live-cell fluorescence microscopy data

**DOI:** 10.64898/2026.05.06.723263

**Authors:** Colin Hogendorn, Ingrid R. Aragon, Samuel Dallon, Eric Batchelor

**Affiliations:** Department of Integrative Biology & Physiology, University of Minnesota Medical School – Twin Cities, Minneapolis, MN, 55455; Masonic Cancer Center, University of Minnesota Medical School – Twin Cities, Minneapolis, MN, 55455

## Abstract

To properly respond to their environment, cells adjust the activity of key regulatory proteins and rates of gene expression. Methods to detect and quantify these forms of regulatory dynamics in living cells are of central importance for understanding cellular signaling events in both physiological and pathological conditions. Current technologies in this field make use of fluorescent probes to track cell signaling dynamics. Although these technologies have been used for decades, challenges remain. In particular, the segmentation, tracking, and interpretation of single cell dynamic data are time-consuming, prone to subjective errors, and often lacking in standardization across experiments. Here, we present SPIFEE, a data pipeline that uses experiment-dependent parameters to smooth noise and quantify key features of fluorescence data from time-lapse imaging studies. Processing data in this manner enhances and accelerates quantification of live-cell gene and protein expression, simplifies data analysis, and facilitates hypothesis generation.

**Author Summary:** Cells adjust protein activity and gene expression levels over time to respond to changes in their environment, a process referred to as cell signaling dynamics. Quantifying cell signaling dynamics in living cells often uses fluorescent probes, such as green fluorescent protein (GFP) and its spectral variants, to track changes in gene expression or protein activity over time. Challenges inherent in analyzing fluorescence data from single cells stem from biological and experimental noise, time-consuming quantification, and subjective errors. To address these challenges, we developed a computational tool called Signal Processing and Integrated Feature Extraction (SPIFEE). The pipeline improves the quality of fluorescence data analysis by reducing noise and extracting signal features in a way that is both intuitive and objective. The pipeline provides more accurate, rapid, and unbiased quantification of time-lapse microscopy data.

## 1. Introduction

Cells continuously sense and respond to their environment through adjustments of the activities of key regulatory proteins and modulation of gene expression levels^1–3^. Recent studies have shown that cells can encode and decode information through control of the temporal dynamics of the signaling pathways and gene regulatory networks^4–7^. Dynamic features such as timing, duration, and frequency of signaling activity have been shown to have major effects on downstream cellular functions^4^. Many important regulatory systems, including those controlling metabolism, cell growth, cell fate, and stress responses, rely on dynamic patterns such as oscillations or pulses to coordinate appropriate outcomes^8–13^. Well-studied examples include the ERK pathway, which exhibits frequency-modulated activation controlling proliferation and differentiation^11,12^; the NF-κB pathway, which transmits information through fold-change detection mechanisms^10,13^; and the transcription factor FOXO1, whose intracellular localization changes dynamically in response to insulin and oxidative stress stimuli^14^. Another key example is the p53 network, a master regulator of DNA damage and stress response, which shows differential temporal behaviors such as single versus repeated pulses depending on the nature and extent of cellular stress^15–18^.

Understanding these regulatory dynamics is central to deciphering cellular signaling processes. Time-lapse fluorescence microscopy using fluorescent biosensors enables quantification of temporal changes in protein localization and concentration^19^. Although this technology has been indispensable for quantifying signaling dynamics with high temporal resolution and in individual cells, several analytical challenges persist. Segmentation, tracking, and feature assignment of single-cell trajectories are often noisy, labor-intensive, and prone to subjective error.

Fluorescence time-series analysis presents significant challenges due to noise arising from both biological variability and experimental measurement, necessitating the use of signal processing techniques such as smoothing and filtering to recover meaningful dynamics^20,21^. In parallel, peak detection and feature extraction methods have been widely developed to identify structured events within noisy biological signals^22,23^. However, these approaches have largely been developed for highly specialized or fully automated peak-detection workflows, rather than for settings that require transparent parameter tuning and interpretability by end users. In this work, we instead focus on developing a framework that is intuitive and accessible to non-experts. While existing computational tools address aspects of fluorescence imaging and time-series analysis^24–26^ they are typically implemented as end-to-end analysis pipelines that prioritize automation. Consequently, they offer limited flexibility for standardized, reproducible parameterization across heterogeneous fluorescence datasets.

However, these approaches have largely been developed for highly specialized or fully automated peak-detection workflows, rather than for settings that require transparent parameter tuning and interpretability by end users. In this work, we instead focus on developing a framework that is intuitive and accessible to non-experts, enabling users to understand, adjust, and reproduce key analysis parameters across datasets. While existing computational tools address aspects of fluorescence imaging and time-series analysis^24–26^, they are typically implemented as end-to-end pipelines that prioritize automation. Consequently, they offer limited flexibility for standardized, reproducible parameterization across heterogeneous fluorescence datasets.

Here we present a pipeline to address some of these challenges -- the Signal Processing and Integrated FEature Extraction (SPIFEE). SPIFEE is a computational pipeline with a graphical user interface implemented in MATLAB that uses raw fluorescence data as an input, uses experiment-dependent parameters to smooth out noise, and yields visually intuitive and informative features (Fig. 1). MATLAB was chosen to preserve consistency with previously published workflows^27^. While SPIFEE does not attempt to impose an absolute fluorescence scale or correct for instrument specific variability, it is designed to work within these constraints. The pipeline standardizes filtering and smoothing procedures to reduce noise and improve internal consistency within time-lapse datasets, thereby enhancing the robustness and reproducibility of downstream quantitative analyses.

**Fig. 1.**
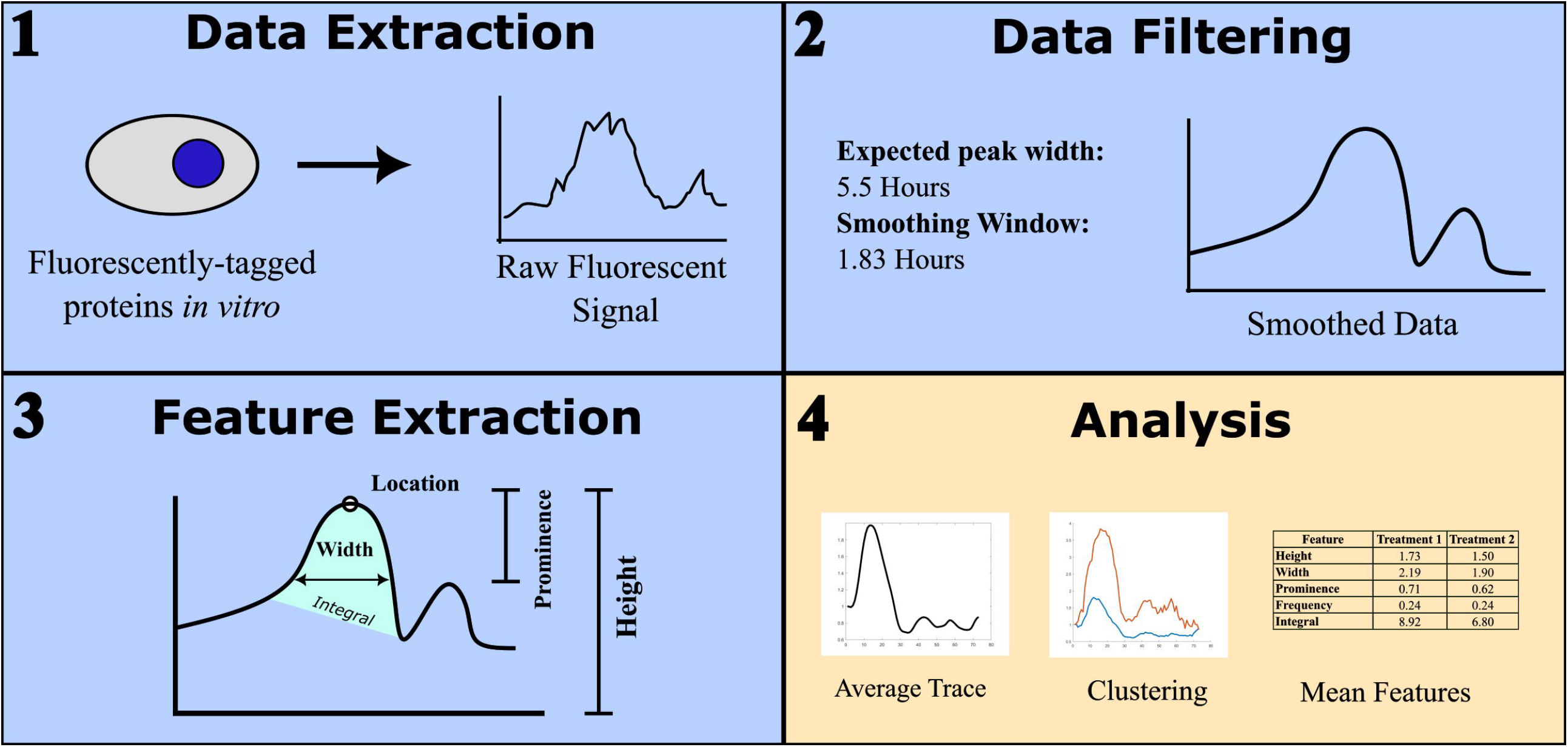
SPIFEE pipeline overview. Data is in the form of time-series traces. Data is first inputted as raw traces, then goes through a series of filtering steps, and features and characteristics of the peaks are then extracted from filtered traces, and then downstream analysis and plotting is performed.

SPIFEE implements standardized feature extraction. Our pipeline introduces a “Default” parameterization set that derives peak-detection constraints from fundamental properties of the dataset and passes them to MATLAB’s *findpeaks*() function. Although fluorescence time-series measurements are often noisy due to both biological and experimental sources, many signaling pathways exhibit characteristic temporal structures that can serve as internal reference points. For example, in response to DNA double strand breaks the intracellular concentration of p53 oscillates with a period of ∼5.5 hours^17^, providing a biologically motivated constraint on peak spacing. SPIFEE uses experimental parameters such as experiment duration and average signal amplitude to inform the *findpeaks*() thresholds that distinguish true biological events from background fluctuations (Table 1). By promoting consistent feature quantification, SPIFEE enhances reproducibility and comparability across studies. Users can also tailor parameters to specific experimental conditions, with all modifications automatically recorded within the SPIFEE output. This process ensures transparency and provides standardization.

**Table 1.** Equations defining *findpeaks()* peak detection parameters. Thresholds for peak height and prominence are scaled to the mean maximal signal across traces, while peak width and spacing constraints are normalized to sampling density and oscillation frequency (in hours).

| ***Parameter*** | ***Definition*** |
| --- | --- |
| $minHeight = \frac{mean(\max(traces))}{10}$ | The minimum height of a peak is 1/10 the mean of the max of every trace. |
| $minProminence = \frac{mean(\max(traces))}{20}$ | The minimum prominence of a peak is 1/20 the mean of the max of every trace. |
| $minWidth/Distance = \frac{\frac{Points}{Hours} * Frequency}{10}$ | The minimum distance between peaks and the minimum width of a peak is a tenth of the pulse period of peaks in terms of Hours. |
| $maxWidth = \frac{Points}{Hours} * Frequency * 10$ | The maximum width of a peak of a peak is ten times the pulse period of peaks in terms of Hours |

## 2. Design & Implementation

### 2.1 Overview

SPIFEE is comprised of a series of scripts and an accompanying graphical user interface (GUI) in the MATLAB language. SPIFEE was specifically developed to analyze oscillatory fluorescence data and extract key features from single-cell traces, although it can effectively analyze other forms of time series data. It uses a combination of MATLAB’s Signal Processing Toolbox along with the Statistics and Machine Learning Toolbox. Users provide two key experimental parameters: the total experiment duration and the average expected oscillation frequency of the measured signal.

Installation requires downloading SPIFEE and adding the resulting folder to the path file in MATLAB. The GUI is launched with the command “SPIFEE_GUI”. Users are prompted to select the fluorescent trace file(s) to be analyzed. SPIFEE natively handles single file or multiple file inputs (Fig. 2). The preferred input data format is a .mat or .csv file where each column represents an individual cell trace. SPIFEE automatically verifies input fluorescence data are provided in the correct matrix orientation by computing a jaggedness metric for each possible orientation. (Supplemental text, Equations S1-S2, and Fig. S1). Incorrectly oriented data introduce artificial discontinuities by alternating between distinct cell traces at successive time points, leading to substantially increased jaggedness. While user input is not overridden, SPIFEE issues a warning when the imputed orientation is likely incorrect. Along with the initial required inputs, users have the option to filter missing values. By default, SPIFEE excludes traces with more than 10% missing data points. Users may change this threshold and optionally normalize traces by either their maximum or initial value. (Supplemental text and Figs. S2, S3). SPIFEE also provides downstream analysis and visualization options, including clustering with multiple distance metrics, per-condition averaged trace plots, heatmaps, and summary statistics of extracted peaks. While these downstream analyses are not intended to be exhaustive, the structured SPIFEE output is designed to facilitate further customized analyses and visualization by users.

**Fig. 2.**
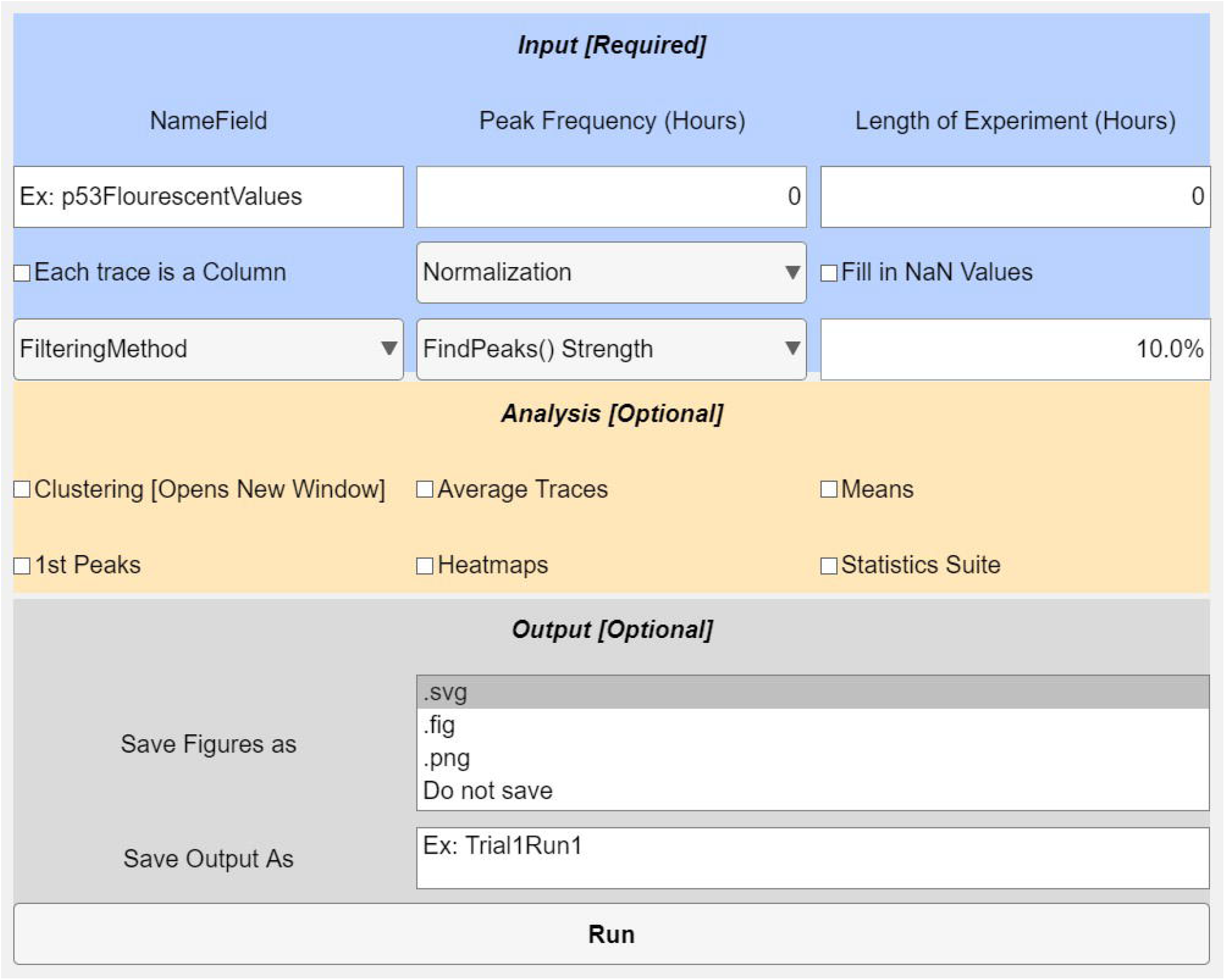
SPIFEE graphical user interface (GUI). The interface is organized into three sections: Input, Analysis, and Output. The Input panel allows users to define experimental parameters, including experiment duration, oscillation frequency, data field location, trace orientation, normalization method, and handling of missing values. The optional Analysis panel is used for selection of downstream visualizations and analyses (e.g., clustering, average traces, first-peak metrics, and heatmaps). The Output panel specifies file naming and data saving options. The analysis pipeline is executed upon selecting Run.

SPIFEE generates a plain English reproducibility record for each analysis run. This output provides a complete description of all user-specified inputs, derived parameters, MATLAB version, and execution metadata required to reproduce a given analysis. It documents how user defined temporal inputs are converted into smoothing window parameters and how these values are turned into peak-finding constraints along with any other important data handling consideration. This plain English record is saved as a text file and is also stored in the SPIFEE output structure (Supplemental text and Fig. S4).

### 2.2 Data Smoothing and Quality Control

To accommodate a range of user needs, SPIFEE offers multiple options for data normalization, smoothing, and imputation. Data may be normalized by either their initial value or maximum value. Following normalization, noise filtering and data smoothing are applied by default using a Gaussian window with a width equal to one third of the expected pulse period. SPIFEE offers other data smoothing methods of various strengths (Supplemental text and Fig. S5). To ensure robustness across experiments of varying durations and pulse dynamics, the smoothing window is adaptively scaled based on total experiment length and the expected peak duration of the target signal (Fig. 3). Adaptive Gaussian or Savitsky-Golay smoothing in SPIFEE is intended for proper feature extraction and analysis of peaks within traces. It is important to note that data with insufficient temporal resolution may not adequately resolve peak dynamics, independent of smoothing (Supplemental text and Fig. S6). Because smoothing modifies frequency content^28^, downstream user spectral analyses should be performed on unsmoothed or minimally processed traces. SPIFEE by default does not include traces that have over 10% missing values. Users may adjust this threshold in the GUI. Traces with missing data under this threshold have data imputed using linear interpolation from nearest neighbors. All data filtering choices are reflected in the plain English output, with information about which traces were or were not imputed provided in the output structure.

**Fig. 3.**
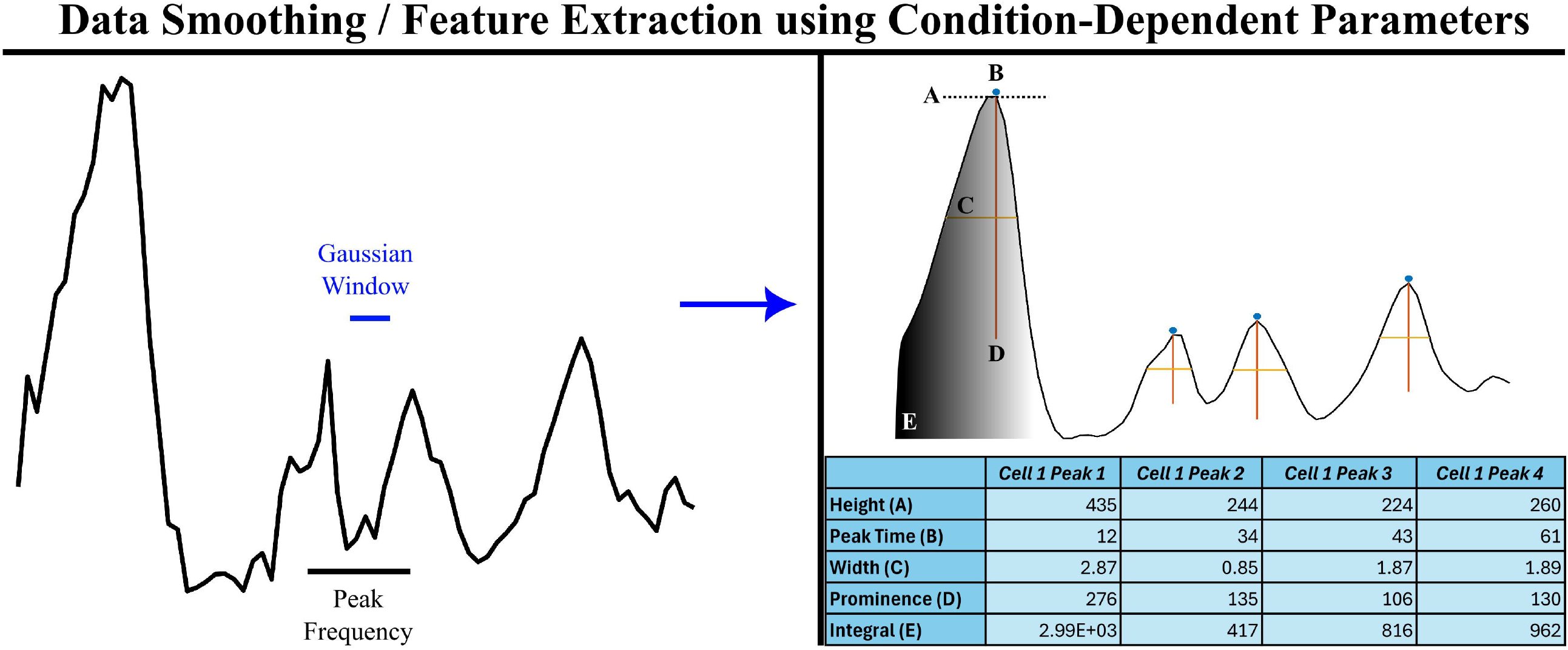
Condition-dependent smoothing & feature extraction of p53 fluorescence traces in MCF7 cells. Fluorescence time series from a 24 h shown before and after Gaussian smoothing. The smoothing window and the expected peak-width timescale are overlaid, demonstrating adaptive parameter scaling based on experiment duration and oscillation frequency. Peaks features are extracted and were identified using the *findpeaks()* function, and the corresponding features of height, location, width, and prominence are annotated.

### 2.3 Pulsatile Dynamics Peak Detection and Extraction

A common feature identified as important in a range of signaling systems is the timing of signal peak activation^8,11^.To identify signaling peaks in an objective manner, once the data is smoothened, peak detection is performed using MATLAB’s *findpeaks()* function with parameters that adapt to the relative scale of the data. We identified five key parameters that optimized peak detection across different experimental contexts (Table 1).

From our survey of dynamic gene expression data, many biologically relevant oscillatory processes occur on the timescale of a few hours^29^. Guided by this range, the parameters presented in Table 1 provide a flexible and efficient approach for peak detection across diverse experimental contexts, offering substantial time savings compared with manual analysis. The pipeline is designed to accelerate data processing while balancing accuracy and computational efficiency, facilitating application to large datasets. Parameters are not universally optimal, and SPIFEE allows users to adjust them to accommodate specific experimental conditions.

After preprocessing, features are extracted from the smoothened data and downstream analysis, including statistical analysis, is performed.

## 3. Results

### 3.1 Extraction of peak features from traces

Pulsatile dynamics, including oscillations, have been discovered as a mode of expression in important signaling regulators including p53 and NF-κb^30^. While the dynamics have been shown to affect cell fate decisions^18^, the specific information encoded within these dynamics to favor specific cell fate responses is less well understood. One feature of these pulsatile dynamics is that across conditions, pulse period remains relatively consistent, making it a powerful anchor point for comparative analysis across experimental conditions. It has been shown that p53 dynamics have a period of around 5.5 hours between pulses^9,16^, which has been reported for p53 as well as many other pulsatile transcription factors^10,31^. Using these ground reference points, we can extract diverse features from the pulsatile biological signals (Figs. 1 and 3).

Processing of each oscillatory peak yields the following information: Height, Location, Width, Prominence, Frequency, and Integral. For example, SPIFEE analysis of an example fluorescence trace from MCF7 cells expressing fluorescently-tagged p53^32^ yielded four peaks with associated features for each peak (Fig. 3).

### 3.2 Extraction of peak features using p53 pulse-controlled data builds ground truth

In developing SPIFEE, our previously published training set was used for validation. The set is derived from a study by Harton *et al*^32^ in which p53 pulses were generated by varying the dosing schedule of the small molecule Nutlin-3, an MDM2 inhibitor. The pharmacological perturbations created six different conditions of p53 oscillations: low amplitude, high amplitude, low frequency, high frequency, long duration, and natural oscillation p53 pulse characteristics (Fig. 4). The resulting data sets served as ground truth for SPIFEE analysis of fluorescence trace peak characteristics.

**Fig. 4.**
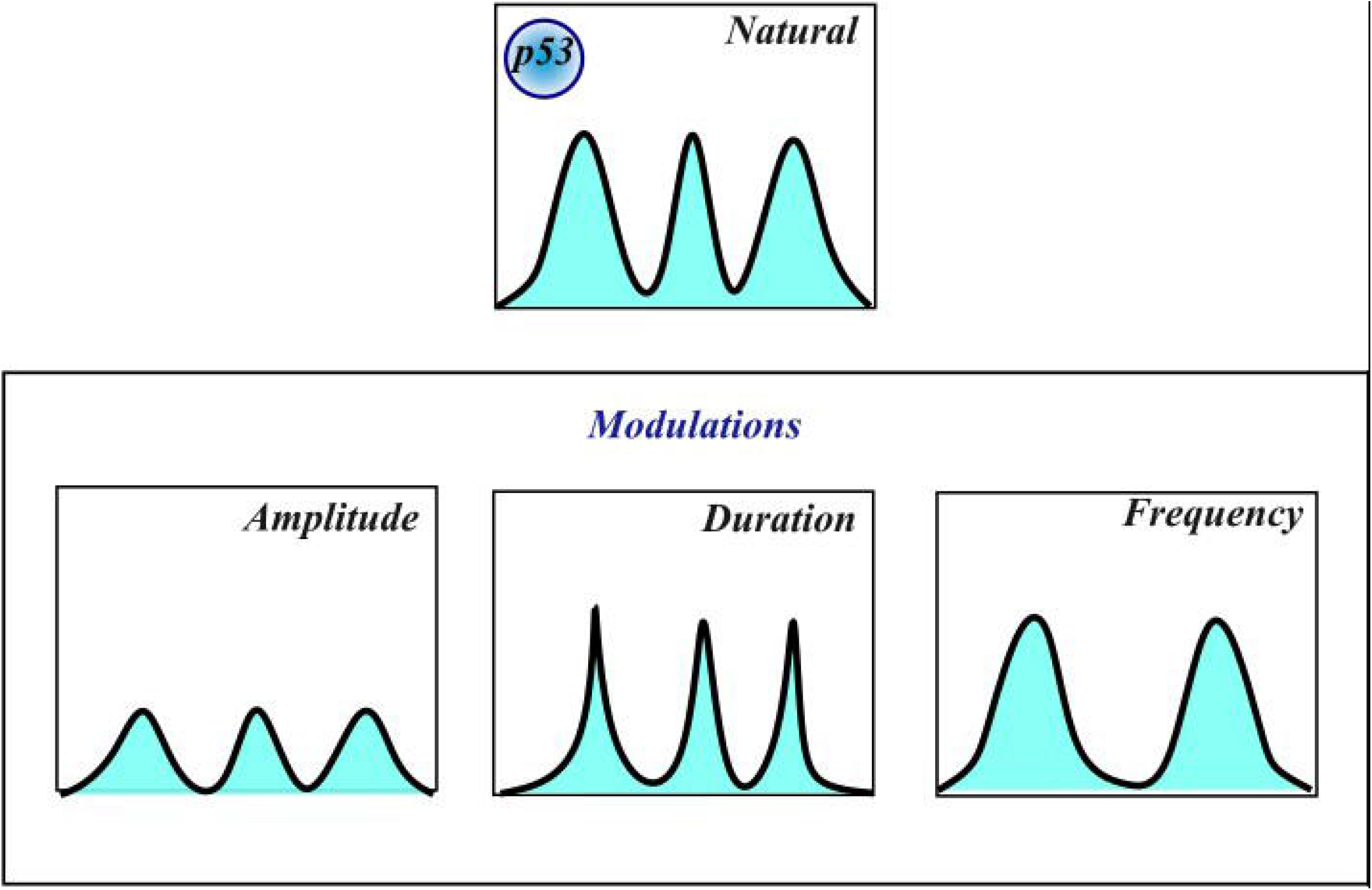
Schematic of the experimental setup for generating manually modulated p53 pulses. The dataset was used to develop SPIFEE. Modulations were performed with Nutlin-3, encompassing six conditions: High Amplitude, Low Amplitude, High Frequency, Low Frequency, Long Duration, and Natural.

### 3.3 SPIFEE Results reinforce ground truth findings

SPIFEE analysis of this dataset successfully recapitulated the intended modulation patterns. Mean values of extracted features broadly aligned with expectations (Table 2). For the long-duration condition, peaks averaged approximately 2.14 hours in width, more than an hour longer than in any other condition. Consistent with this manipulation, the long-duration dataset was statistically significantly different from the others in peak width. Using Cliff’s delta, we found an effect size of (0.45). We also found a similarly large effect size in peak integral. The natural condition exhibited a modest negative amplitude effect size (–0.2878), reflecting that the other experimental conditions explicitly manipulated peak amplitude. Apparent discrepancies such as the High Amplitude condition exhibiting a similar average amplitude to the Low Amplitude condition (403.15 and 485.64 AU respectively) reflect inherent heterogeneity in single-cell responses, even under controlled, manually modulated input conditions. These results highlight the complex interdependence of dynamic features. For example, extended pulse duration may inversely affect frequency, while amplitude inherently influences peak integral and related metrics. Despite feature interdependencies, biological noise, and experimental sources of error, the controlled dataset provided a valuable testbed for the development and validation of SPIFEE’s feature extraction. We next assessed the pipeline’s generalizability using independently generated data encompassing distinct biological targets and experimental designs.

**Table 2.** Extracted features of p53 pulses. Experimental conditions from Harton et al set up a clear distinction between features extracted from Nutlin-modulated pulses of p53.

| p53 Manual Modulation Peak Averages |  |  |  |  |  |  |
| --- | --- | --- | --- | --- | --- | --- |
|  | Low Amplitude | High Amplitude | Low Frequency | High Frequency | Natural | Long Duration |
| **Amplitude** | 485.64 | 403.15 | 533.28 | 314.93 | 358.35 | 482.24 |
| **Width** | 35.87 | 34.22 | 34.63 | 1.48 | 1.44 | 2.15 |
| **Prominence** | 237.63 | 171.58 | 344.43 | 144.84 | 136.52 | 223.14 |
| **Frequency** | 0.265 | 0.294 | 0.260 | 0.287 | 0.296 | 0.298 |
| **Integral** | 1798 | 1294 | 1961 | 1015 | 1090 | 2710 |

### 3.4 SPIFEE’s automated peak finding agrees with manual annotation with improved speed

To evaluate SPIFEE’s ability to accurately identify relevant features from fluorescence traces, we performed manual annotation of a dataset and compared these results with features extracted by SPIFEE. To benchmark performance against inter-user variability, we selected 25 traces that were independently annotated by multiple users. Across these shared traces, annotators identified an average of 180 peaks (s.d. = 19). SPIFEE detected 167 peaks on the same traces, falling within the range of manual variation (Fig. 5). The distribution of SPIFEE peaks similarly overlapped with the range observed across annotators, indicating that automated performance is consistent with expert curation We further quantified agreement using the intraclass correlation coefficient (ICC = 0.8785). On a per-user basis, SPIFEE’s peak identification matched manual annotations with an average agreement of 96.15%, demonstrating that the algorithm reliably captures features identified by experts.

**Fig. 5.**
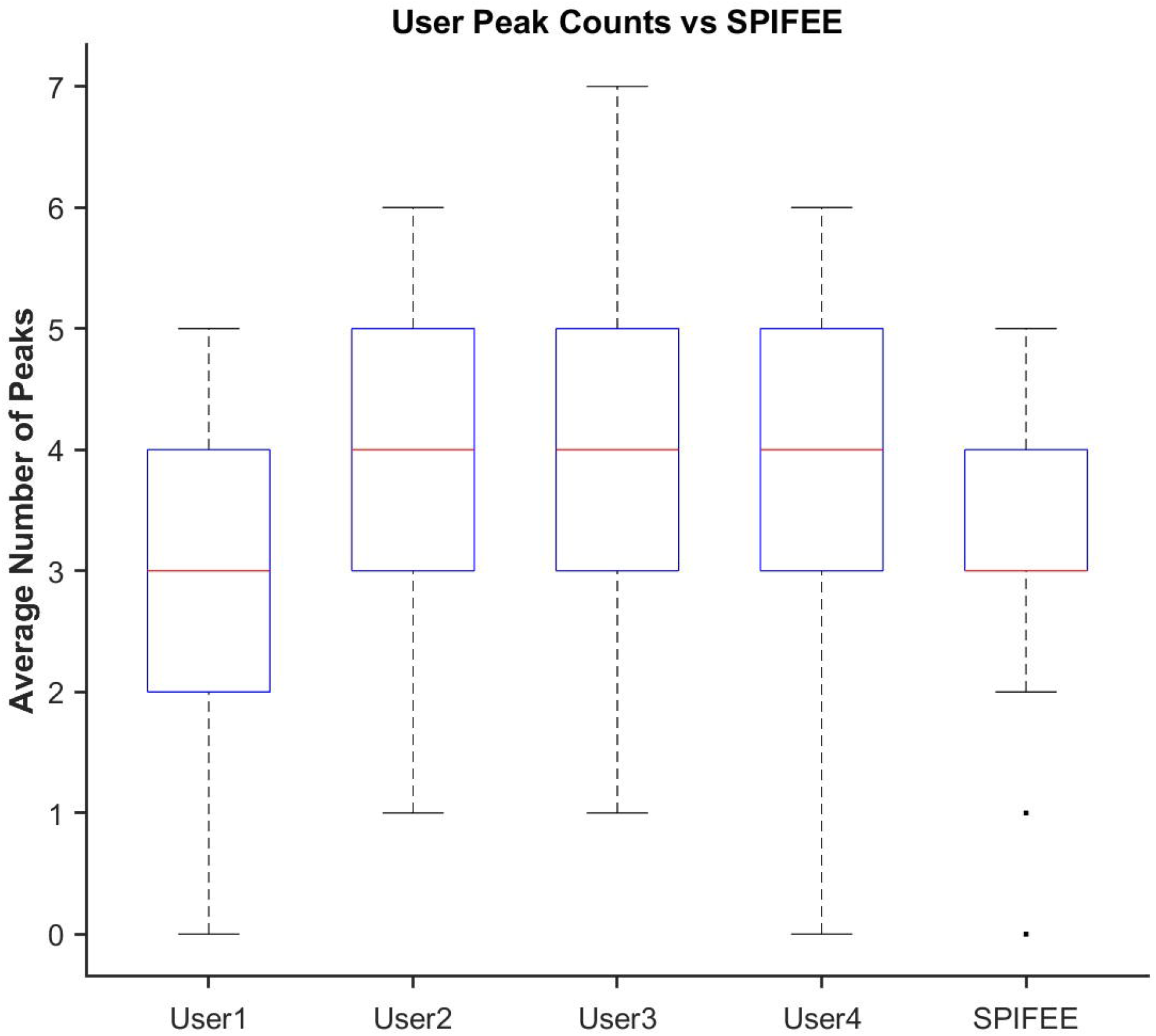
Comparison of manual annotations to SPIFEE annotation for peak finding. Box plots showing the number of peaks identified in 25 shared p53 fluorescence traces by four human annotators and by SPIFEE. Boxes represent the interquartile range (IQR), whiskers denote 1.5× IQR, and center lines indicate medians. SPIFEE’s peak counts fall within the natural variation observed across annotators, demonstrating concordance between automated and manual curation.

Manual annotation was also substantially slower than SPIFEE. To generate the manual annotation set, users were tasked with annotating 50 traces each. Even with a semi-automated workflow to accelerate the process, users required an average of 2 hours and 15 minutes to complete the full set. In contrast, SPIFEE processed the same data in 19.2 seconds (Fig. 6), representing a ∼421-fold reduction in analysis time.

**Fig. 6.**
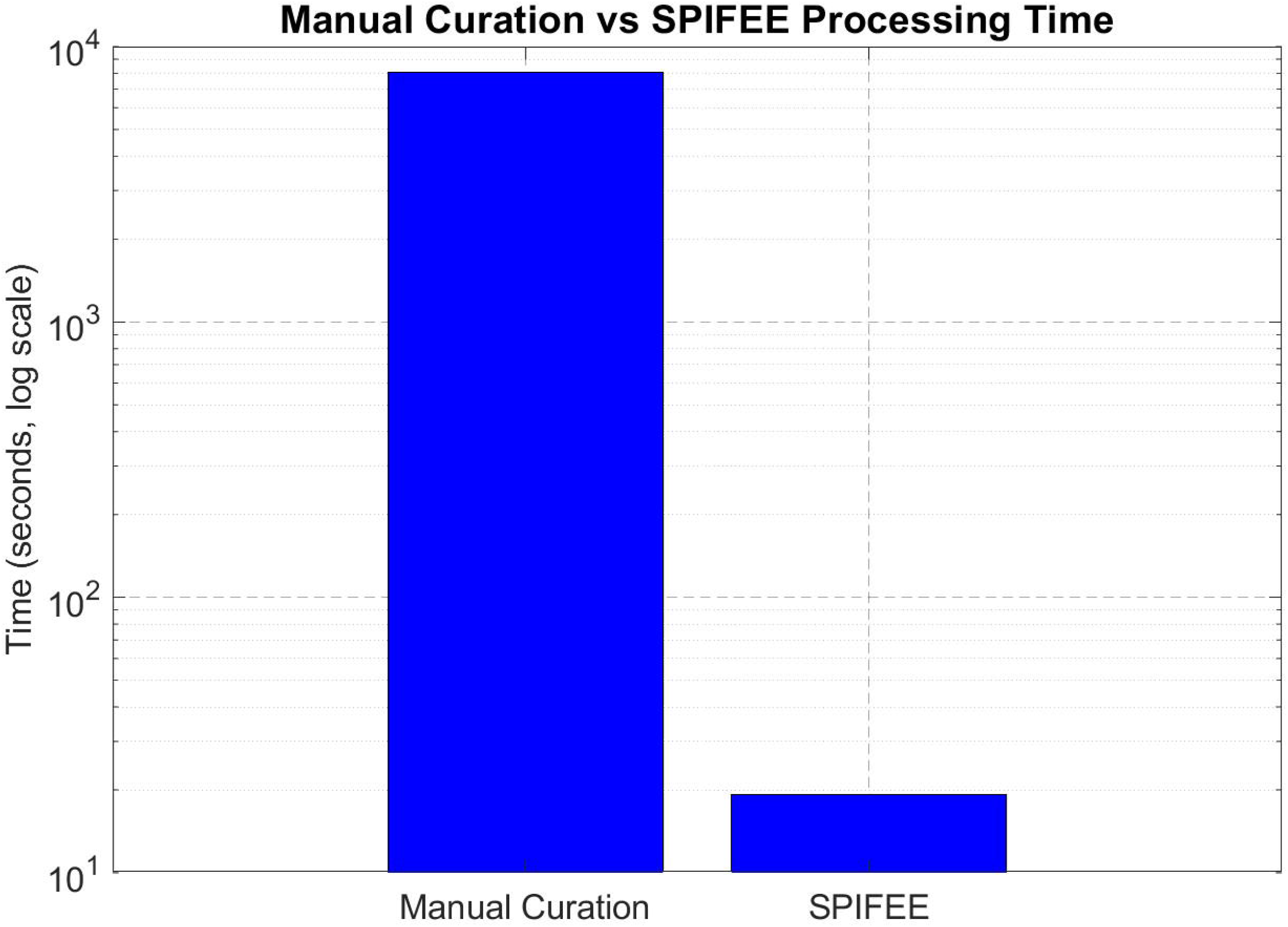
Comparison of the timing of manual annotation vs. SPIFEE annotation. Log scale comparison of Manual annotation of identifying peaks in time series data compared to our pipeline SPIFEE.

We performed the same manual annotation procedure on the p53 modulation dataset to further evaluate the robustness of the peak detection pipeline. Using the default parameter configuration, the method achieved an F1 score of 0.7674, which was the highest performance among the tested filtering windows.

### 3.5 Extraction of peak features from FOXO1 dataset shows SPIFEE generalizability

To evaluate SPIFEE’s generalizabiity, we analyzed a published dataset from an independent laboratory that quantified FOXO1 and p53 expression in response to H_2_O_2_ (Figure. 7) ^14^. Notably, the parental cell line used was the same as in the Harton *et al*.^32^ data set, allowing us to robustly test the performance of our data pipeline in the context of a shared cell line but under different laboratory and experimental conditions. SPIFEE recapitulated the signal processing features extracted and conclusions drawn by the original authors.

**Fig. 7.**
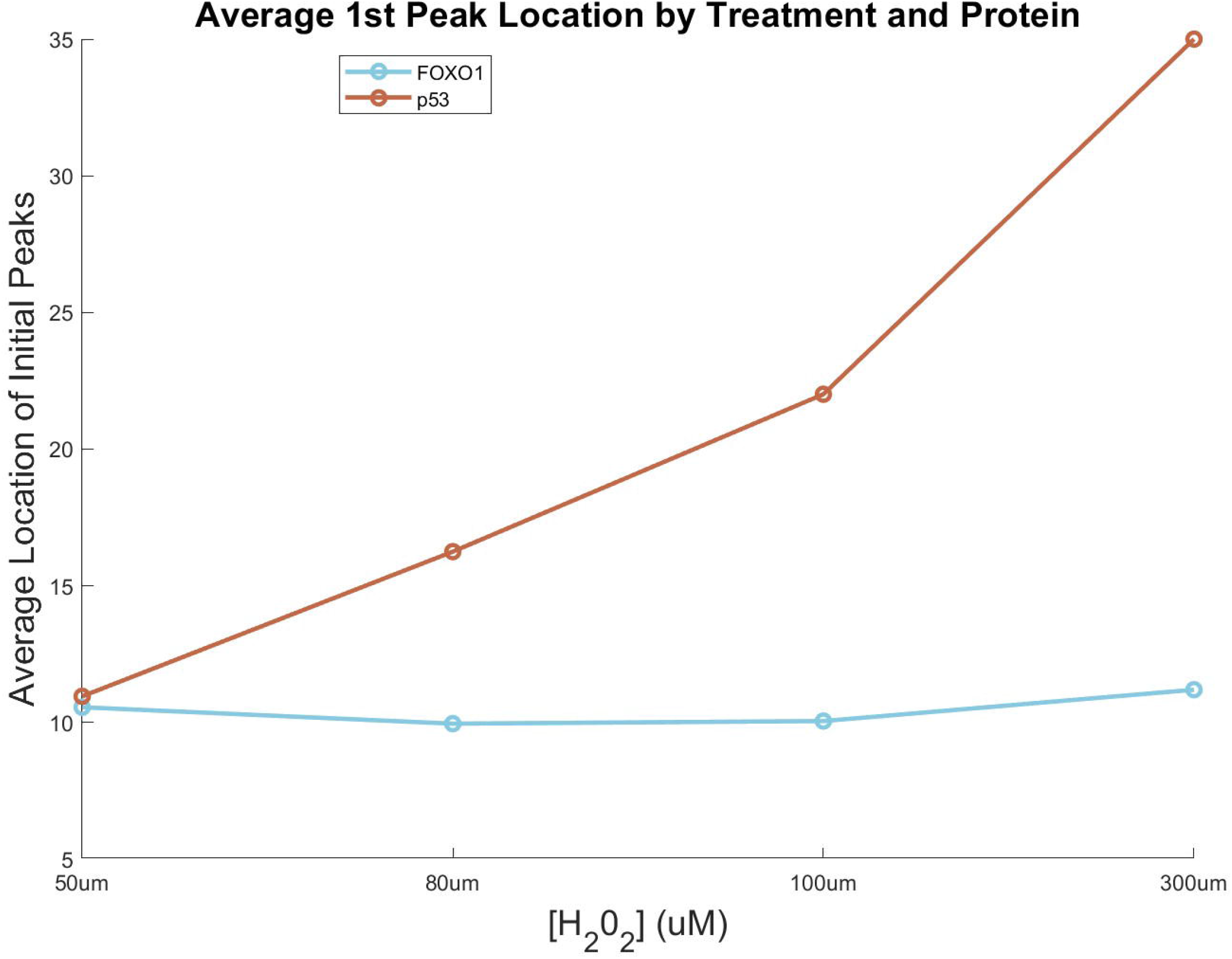
SPIFEE-determined differential responses to H_2_O_2_ for both FOXO1 and p53. Plot of the average location of the first pulse of p53 and FOXO1 in response to increasing H_2_O_2_.

As an example, we quantified the timing of the first peak of p53 and FOXO1. The original study showed a growing delay in the first pulse of the first peak of p53 with higher H_2_O_2_ concentrations, whereas FOXO1 ocurred at relatvitely consistent times. SPIFEE reproduced these findings (Figure. 8). Specifically, the first peak of FOXO1 occurred on average at 10.39 hours across conditions, whereas the initial first peak of p53 ocurred at 10.92, 16.23, 21.72, and 25 hours, respectively, which supported the authors’ findings that as H_2_O_2_ damage increased, p53 expression became more delayed.

**Fig. 8.**
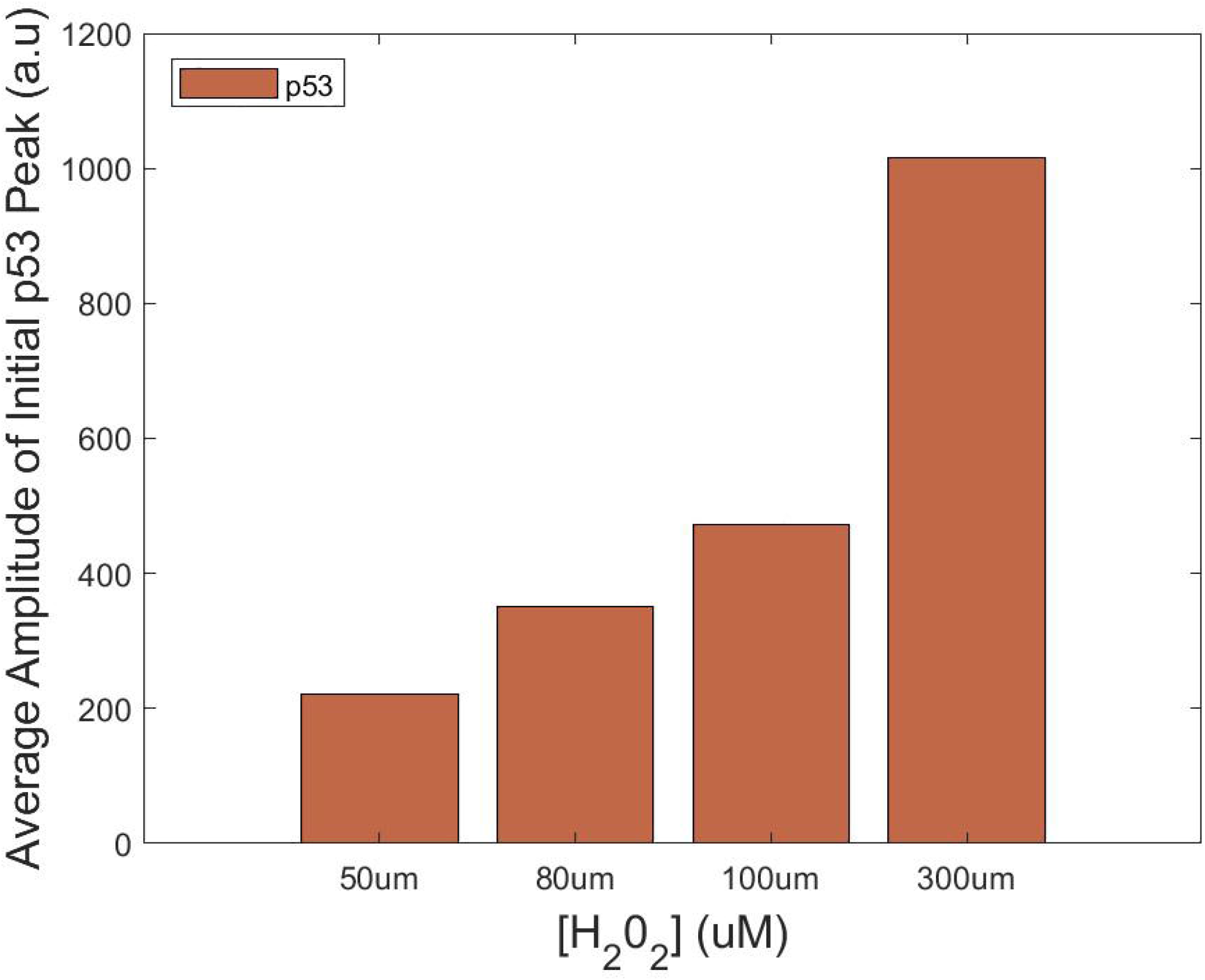
SPIFEE-determined p53 peak amplitude of data from an independent lab. Dose-dependent changes in the first p53 pulse following H_2_O_2_ treatment. The relative amplitude of the first peak increases with higher H_2_O_2_ concentrations, and the timing of this peak exhibits a delay consistent with previously published dynamics of p53 and FOXO1 oscillatory responses.

These results demonstrate that SPIFEE is capable of recapitulating both global and fine-grained dynamic behaviors. In their original analysis, Jose et al^14^ noted, “The dynamics of p53 accumulation also differ in response to higher concentrations of H_2_O_2_. In some cells, p53 levels oscillate similar to the 50 µM dose, yet often with a higher initial spike in p53 levels.” This observation is supported by features extracted through SPIFEE (Fig. 8), which showed increasingly large amplitudes in the first peak of p53 traces in response to high oxidative stress. However, this effect can be obscured by data normalization, underscoring the importance of appropriate preprocessing. Interpretation of SPIFEE outputs must therefore consider both biological and experimental context. For instance, in this dataset, p53 is quantified in arbitrary fluorescence units, whereas FOXO1 is measured as a nuclear-to-cytoplasmic ratio and will have distinct signal ranges and properties.

Although SPIFEE was designed primarily to analyze oscillatory signals in experimental contexts such as p53 dynamics in response to DNA double strand breaks, it also performed effectively on signals with transient burst profiles. FOXO1 nuclear accumulation exhibits multiple dynamic profiles, including sigmoidal shapes and transient bursts in expression^33^. Our pipeline successfully identified and extracted features from these non-oscillatory, transient responses. When evaluated against manually annotated FOXO1 data, SPIFEE achieved an F1 score of 0.6742 with a mean peak height difference of 0.0059 relative to user annotations.

Together, these results demonstrate the utility of SPIFEE diverse signaling contexts. This is reinforced by that fact that although signaling dynamics are broadly put into categories, these categories are not always clearly distinct and depend strongly upon context. A recent review of ERK dynamics stated that it can be grouped into “several major categories, including sustained, transient, peak with sustain, oscillatory, sporadic, and complex”^34^. SPIFEE provides a means to obtain information from a variety of data types and subsequently generate potentially useful visualizations and stores data in an easy to read and use MATLAB data structure. Additional downstream visualizations, graphs and analysis shown are discussed in detail in the Supplementary Text and Figs. S7-S11.

## 4. Discussion and Availability

The SPIFEE pipeline provides a novel workflow for hypothesis generation and rapid analysis of single-cell fluorescence traces through intuitive feature extraction and a GUI. To our knowledge, SPIFEE is the first dedicated pipeline for single-cell fluorescence trace analysis that incorporates adaptive peak filtering to address standardization challenges within the field. By automating complex analyses, SPIFEE improves efficiency and reproducibility, making it well-suited for large datasets and a variety of experimental contexts.

Applications of SPIFEE to publicly available datasets demonstrated its adaptability, versatility, and overall utility. The pipeline streamlines advanced data analysis, making it accessible to researchers with varying levels of computational expertise, while providing a foundation for future developments. Further detailed analysis utilizing SPIFEE output might provide interesting new insights. Since SPIFEE enables rapid analysis, it well suited itself for generating large-scale fluorescence dataset that could support machine learning based discovery approaches. Such efforts may reveal previously unrecognized expression dynamics in key signaling pathway mediators.

## 4.1 Code Availability

The SPIFEE pipeline is an open-source computational tool for processing single-cell fluorescence data. It is available under the MIT License, which permits unrestricted use, modification, and distribution provided proper attribution is given. The pipeline is accessible at GitHub.

https://github.com/ColinHogendorn/SPIFEE

## 4.2 Supplementary Information

Additional figures, equations, and methods are provided: (https://github.com/ColinHogendorn/SPIFEE/blob/main/Supplement_S1.pdf)

## 4.3 Packages and Version control

SPIFEE was developed in MATLAB 2023a and utilizes the Signal Processing Toolbox

## Acknowledgements

We thank Andrew Paek, Myong-Hee Sung, and members of their labs for their help in providing data for the testing of SPIFEE.

This work was supported by R01GM149666 (to E.B.) from the National Institutes of Health.

