## Supplement S1 for "SPIFEE -- A pipeline for analyzing traces of live-cell fluorescence microscopy data"

#### Data Orientation Validation

SPIFEE includes a preprocessing step to validate the orientation of input data matrices (e.g., whether traces are arranged by rows or columns). Incorrect orientation can lead to invalid feature extraction and misleading results. To address this, SPIFEE computes a continuity-based metric for each orientation. The orientation with the higher continuity score is assumed to represent the correct temporal structure of the data. If the inferred orientation differs from the user-specified format, a warning is issued, although the software does not automatically override user input.

For a series of x points:

$$\mathbf{x} = \{x_1, x_2, \dots, x_N\}$$

The **second-difference jaggedness** $J_2$is defined as the mean absolute second finite difference (Equation S1)

$$J_2 = \frac{1}{N-2}\sum_{i=1}^{N-2} | x_{i+2} - 2x_{i+1} + x_i | \quad (N \geq 3) \tag{1}$$

This quantity measures local changes in slope and increases with point-to-point irregularity or discontinuities in the signal.

The **continuity score** $C$is a bounded transformation of $J_2$ (Equation S2):

$$C = \frac{1}{1 + J_2} \tag{2}$$

C will be a value from 0 to 1, with 1 indicating perfect continuity. As a demonstration, we use a rather jagged example trace from a 7 day mcf7 in response to doxorubicin and measure its continuity scores in the 2 orientations. A higher score denotes a more continuous trace.

Here we see that it indicates the row orientation as the better scored orientation which is correct. This demonstrates that even with jagged data, our score is robust with respect to identifying the right orientation of the data. If the orientation identified doesn't align with user input, it will throw a warning but will not forcibly change orientation for the user (Fig. S1).

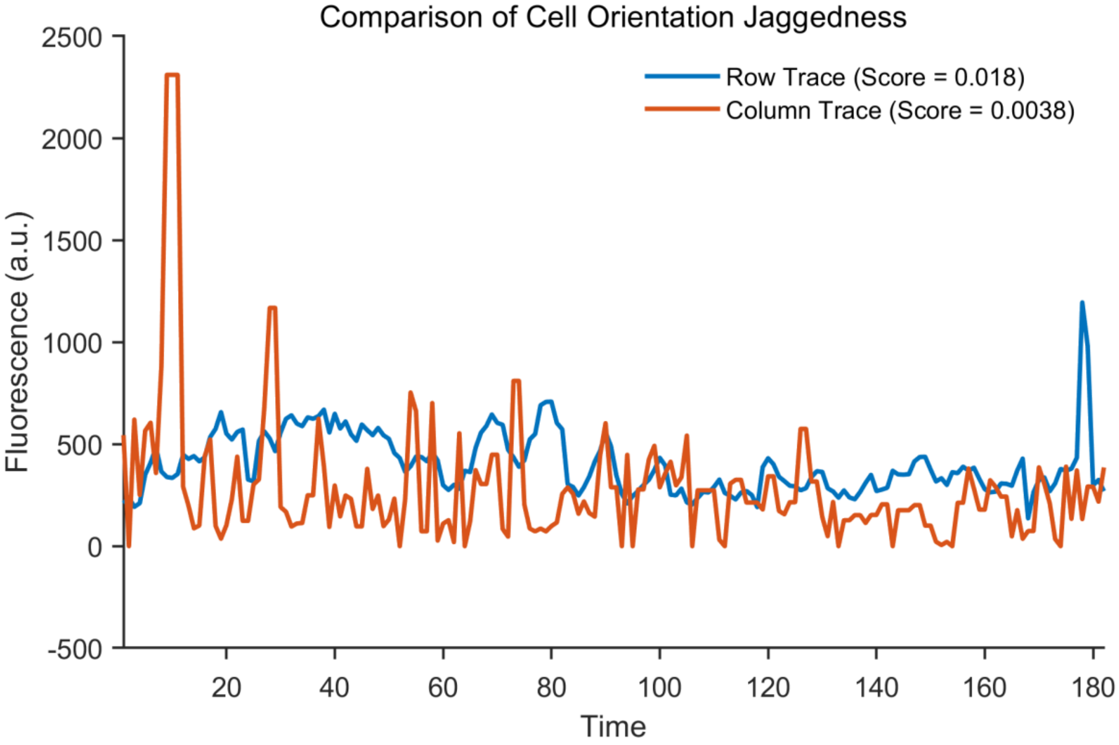

**Fig. S1: Jaggedness scores for a representative fluorescent trace in both orientations**. The correct orientation yields a higher continuity score, reflecting improved preservation of temporal structure. This approach remains robust even for highly variable signals.

#### Missing Values and Quality Control

To start the filtering process, SPIFEE by default, filters out traces that have over 10% of their points missing. If selected, SPIFEE will impute data using linear interpolation from nearest neighbors for traces under that threshold. In the GUI, users may input a higher missing data threshold.

If there are NaN values anywhere in the data, SPIFEE will automatically generate quality control graphs. We calculate pointwise missingness along the time points and then evaluate and plot an "optimal" cut off point (Fig. S2). Note that this cutoff is merely a way of visualizing the missingness of data, and not a key part of the SPIFEE process. We then evaluate and plot the amount of data that passes the NaN threshold or not (Fig. S3). It is important to note that SPIFEE needs traces to be complete to compute downstream feature extraction / scripts.

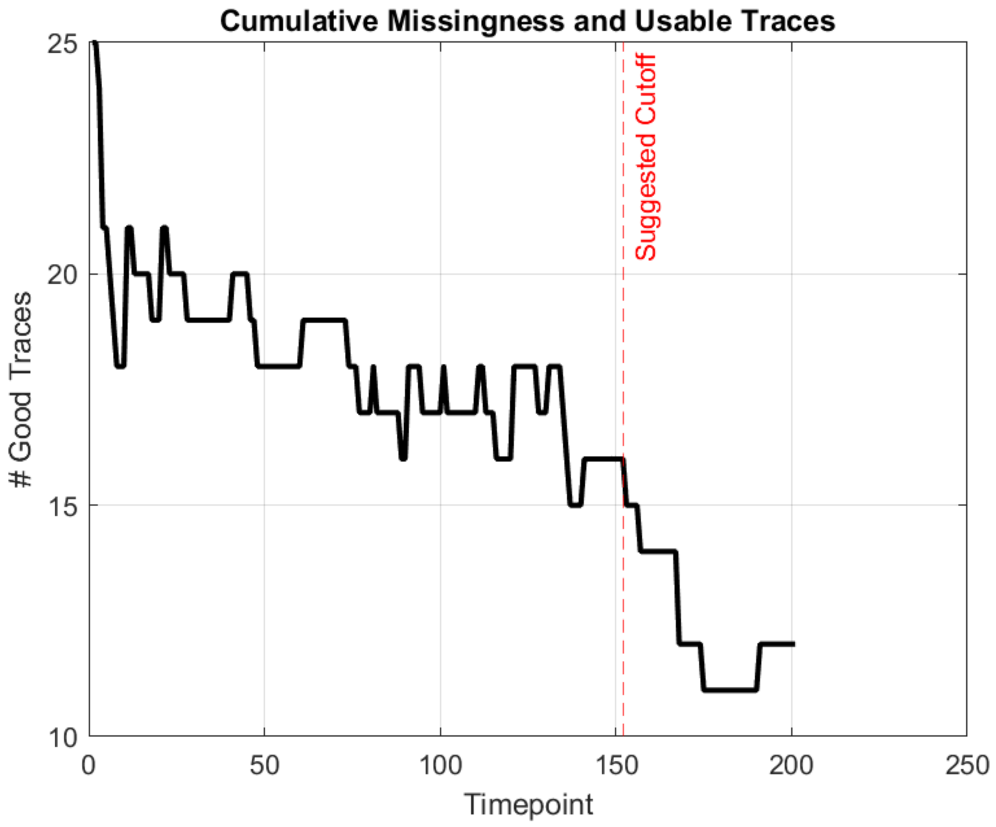

**Fig. S2: Pointwise missingness across traces**, A suggested cutoff is shown that balances data retention with the number of traces included in downstream analysis.

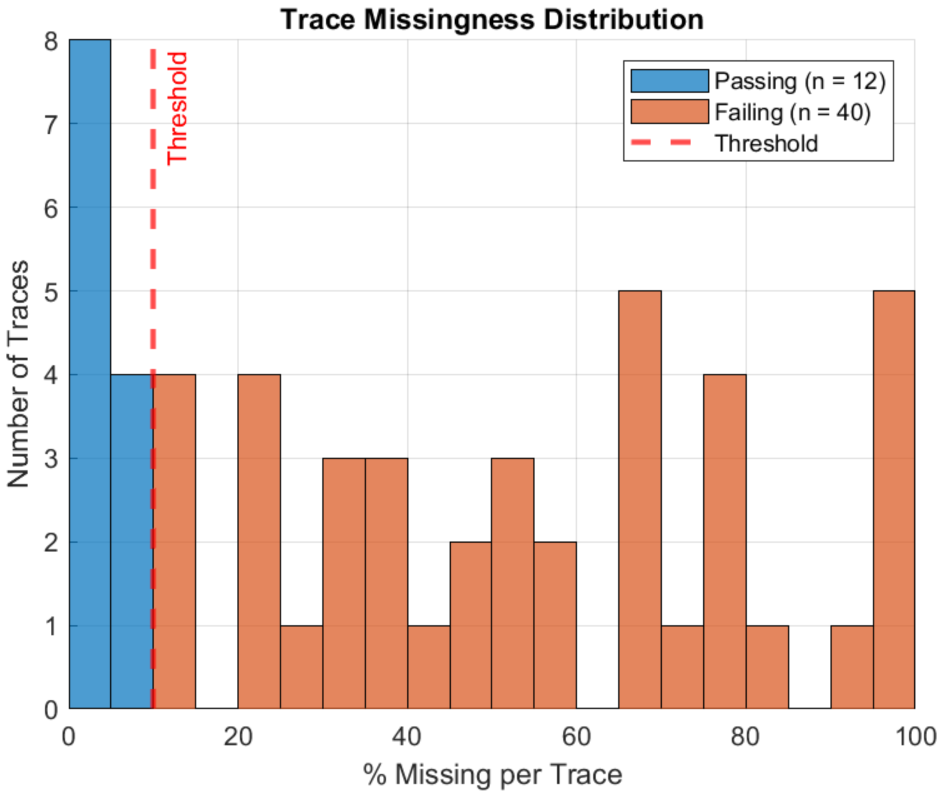

**Fig. S3. Percent missingness per trace.** Data traces have % missing calculated per trace, and are shown passing or failing the % missing data threshold. The % threshold by default in SPIFEE is 10%.

#### Parameter Reproducibility

All user-defined and default parameters within SPIFEE are stored in the output data structure. This includes filtering parameters, peak detection thresholds, clustering settings, and data preprocessing options. All these considerations are put together and reported in a simple "plain English" output. Along with storing within the output data structure, SPIFEE also outputs this plain English output as a text file. In addition to being embedded in the output structure, SPIFEE exports this summary as a standalone text file for ease of sharing and documentation. We strongly recommend reporting this parameter summary alongside all results to support reproducibility. Because feature extraction outcomes can be sensitive to parameter selection, explicit documentation of analysis settings enables direct comparison across experiments (Fig. S4).

```
SPIFEE Analysis Summary
------------------------
Analysis Date and Time:
10-Apr-2026 10:44:54

Software Environment:
9.14.0.2239454 (R2023a) Update 1

Data files used:
 - TreatFiber.mat
 - TreatNone.mat
 - UnTreatFiber.mat
 - UnTreatNone.mat

Preprocessing and Data Handling:
--------------------------------
Input traces were analyzed as being over a user-defined duration of 24 hours.
The Data had 52 Traces with 201 points
Data were smoothed using a Gaussian filter derived from a user-specified oscillation frequency of 5.5 hours.
The resulting smoothing window size was 15.3542 points, corresponding to 1/3rd the length of the user's input frequency.

Data was Not Normalized
Data was inputted as being structured Horizontally
For NaN values, there was data imputation with fillmissing() with linear interpolation from nearest EndValues.
Only traces with less than 10 percent of points missing were imputed.
Traces exceeding the missing-data threshold of 10% of points were excluded from downstream analysis.

Peak Detection Parameters:
--------------------------
Peaks were identified using the following criteria:
Minimum peak height: 37.80162
Minimum peak prominence: 25.20108
Minimum peak-to-peak distance: 0.55 hours
Minimum peak width: 0.55 hours
```

**Fig. S4. SPIFEE Reproducibility Report:** Human-readable summary of all user-defined and default SPIFEE parameters, including filtering, peak detection, clustering, and preprocessing settings. This output is designed to support transparent reporting and reproducibility across analyses.

#### Filtering Methods Overview

We evaluated multiple smoothing approaches at first, including Gaussian filtering, moving average filters, locally estimated scatterplot smoothing (LOESS), and Savitzky–Golay filtering. Each method was assessed based on its ability to reduce noise while preserving biologically relevant peak features.

Moving average filters produced results largely redundant with Gaussian filtering, without offering additional advantages in peak preservation. LOESS filtering introduced spurious local extrema under certain parameterizations, leading to false-positive peak detection. In contrast, both Gaussian and Savitzky–Golay filters provided a favorable balance between noise reduction and feature preservation. (Fig. S5).

Based on these observations, SPIFEE implements Gaussian and Savitzky–Golay filtering as primary smoothing options, with user-adjustable parameters to accommodate variability across datasets.

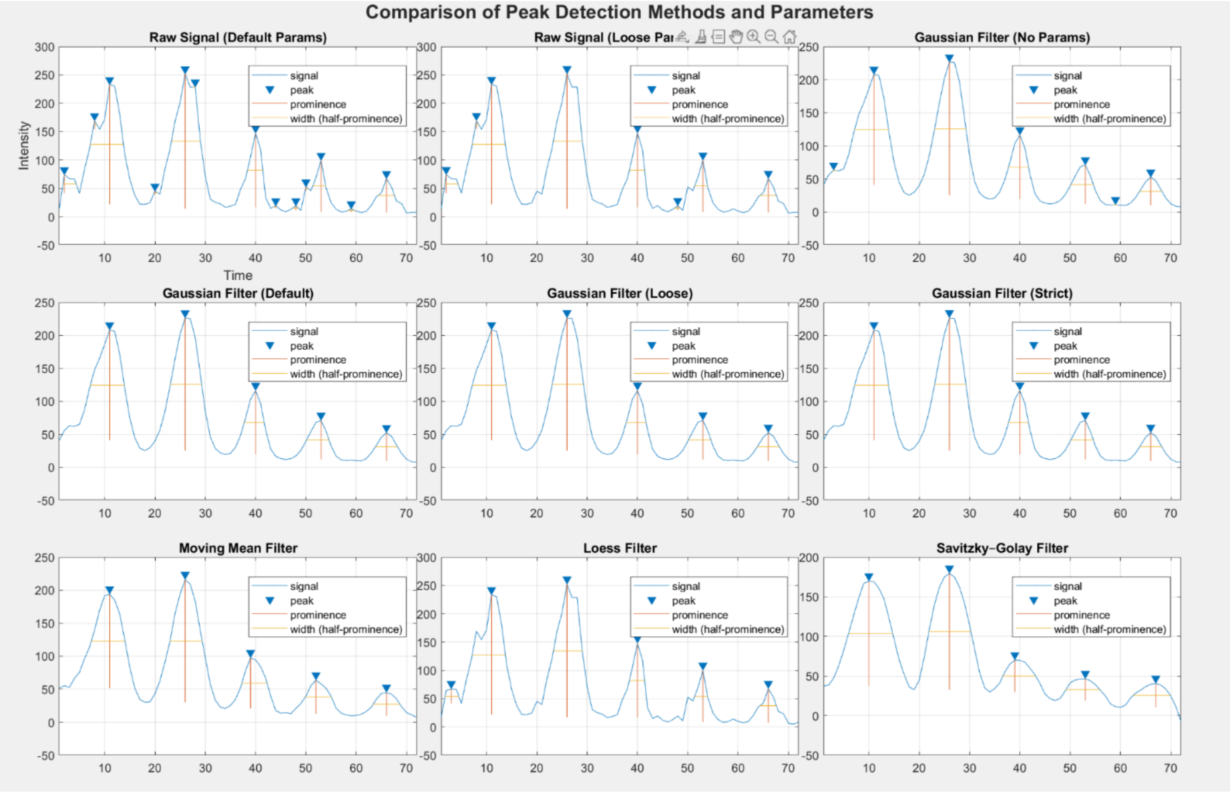

**Fig. S5. Evaluation of smoothing approaches.** Evaluation of multiple smoothing approaches applied to representative fluorescence traces prior to peak detection using findpeaks(). Methods include Gaussian filtering with multiple window sizes, moving average filtering, LOESS smoothing, Savitzky–Golay filtering, and unfiltered traces. Filter performance was assessed based on downstream peak detection consistency and qualitative preservation of signal structure.

To further evaluate the two filtering windows, we compared Gaussian and Savitzky–Golay filters across a range of window sizes. Filtered outputs were processed and benchmarked against a manually annotated p53 dataset. Overall, we observed that the default Gaussian window provided the best balance between precision and recall, yielding the highest F1 score among tested configurations (Table S1). While looser Gaussian settings slightly improved recall, they substantially reduced precision. Conversely, stricter settings performed worse in both metrics. Savitzky–Golay filtering demonstrated competitive performance but had a lower F1 score overall. Gaussian performed the best across every condition in the 6-condition set, demonstrating its ability across contexts.

| Filter Name | True Positive (TP) | False Positive (FP) | False Negative (FN) | Precision | Recall | F1 |
|---|---|---|---|---|---|---|
| *Gaussian Default* | 721 | 194 | 243 | 0.7880 | 0.7479 | 0.7674 |
| *Gaussian (Strict)* | 460 | 135 | 504 | 0.7731 | 0.4772 | 0.5901 |
| *Gaussian (Loose)* | 733 | 514 | 231 | 0.5878 | 0.7604 | 0.6630 |
| *Savitsky-Golay* | 762 | 447 | 202 | 0.6303 | 0.7905 | 0.7013 |
| *Savitsky-Golay (Loose)* | 727 | 561 | 237 | 0.5644 | 0.7541 | 0.6456 |
| *Savitsky-Golay (Strict)* | 592 | 104 | 372 | 0.8506 | 0.6141 | 0.7133 |
| *Unfiltered* | 727 | 561 | 237 | 0.5644 | 0.7541 | 0.6456 |

**Table S1: Comparison of peak detection performance on filtering methods and strengths.** Filtering and parameterizations were compared on a manually curated ground truth dataset. Metrics include true positives (TP), false positives (FP), false negatives (FN), precision, recall, and F1 score. The default Gaussian filter configuration achieved the highest overall F1 score, indicating the best balance between sensitivity and specificity across tested methods.

#### Peak Detection Sensitivity to Smoothing

The ability to detect transient features is fundamentally limited by the temporal resolution of the input data. Peaks can only be identified if they are adequately sampled relative to their duration. If the characteristic width of a peak is smaller than the sampling interval (e.g., points per hour), the signal will be under sampled, and transient events may not be detectable. SPIFEE does not attempt to reconstruct features below the sampling limit. Accordingly, users should ensure that data acquisition frequency is sufficient to resolve the temporal dynamics of interest prior to analysis.

Choice of window length for data smoothing is itself a nontrivial topic and has ramifications on data feature extraction. Although not the primary focus of this work, it has been studied extensively, most notably by T. Lindeberg in the context of scale-space theory(1). A central insight from this framework is that there is no single “true” representation of a signal under smoothing. Instead, meaningful features are those that persist across a range of smoothing scales. Users are therefore encouraged to run SPIFEE with multiple filtering strengths to assess peak persistence across scales. Features that remain stable under increasing smoothing are more likely to reflect underlying biological dynamics, whereas transient features are more consistent with noise. Rather than defining an optimal smoothing scale, we emphasize that the selected filters demonstrate robust agreement with manual annotations across diverse datasets and experimental conditions. To illustrate this behavior, we present a 168-hour time-course of p53 expression following doxorubicin treatment, analyzed using multiple Gaussian window sizes (Fig. S5).

A potential concern with the default parameter set is that shorter time-scale recordings may appear visually smoother than longer-duration datasets. This can motivate attempts to enforce a uniform degree of visual smoothness across experiments. However, we find that such normalization can be detrimental to peak detection performance. In the example shown, default parameters correctly identify biologically relevant peaks at approximately 2 and 36 hours. Increasing the smoothing window to achieve a more uniform visual appearance suppresses these features, leading to them not being found by findpeaks(). This highlights a fundamental tradeoff between visual smoothness and preservation of analytically meaningful signal structure, consistent with scale-space theory. It is important to note that features such as absolute height of a pulse are affected by smoothing windows

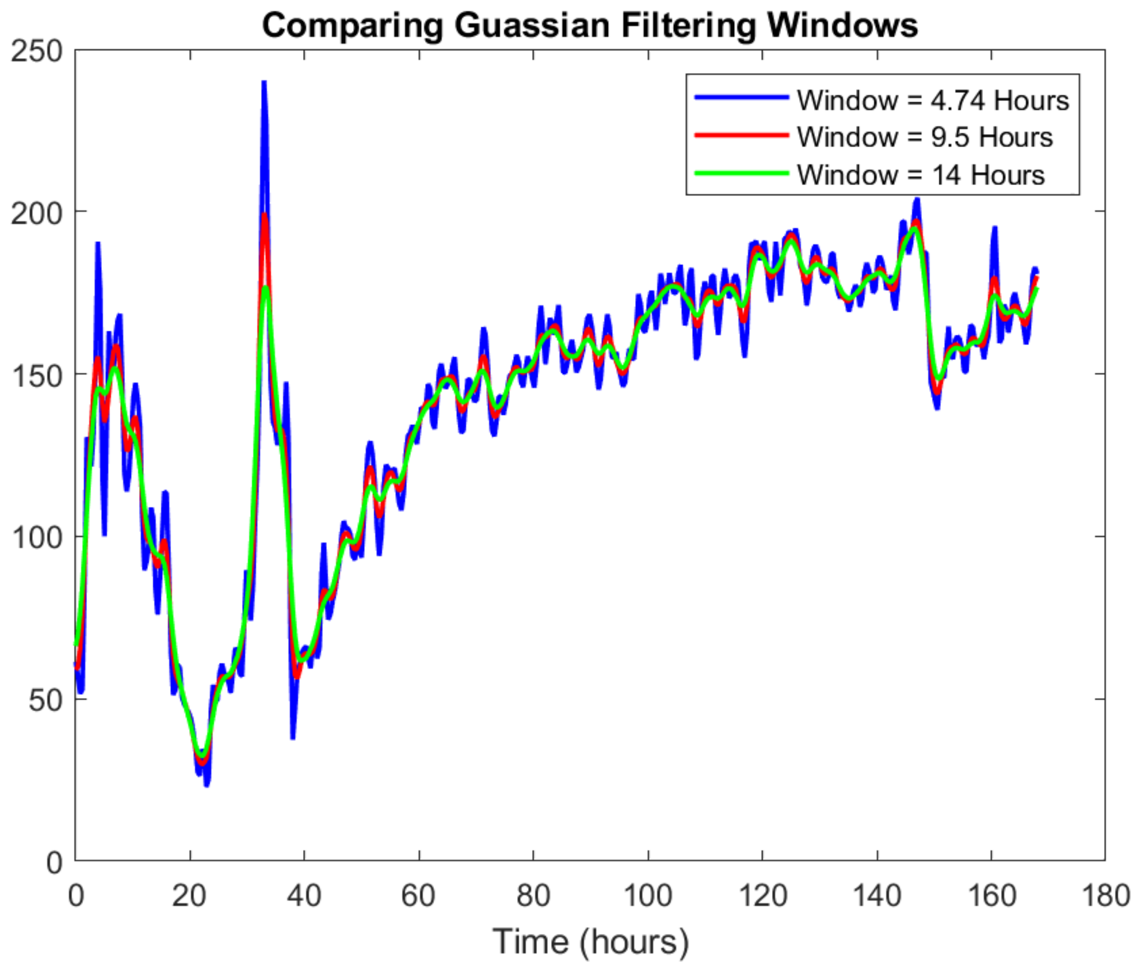

**Fig. S6: Effect of Gaussian smoothing scales on Data**. This representative p53 fluorescence trace taken over 168 hours following doxorubicin treatment, is shown under multiple Gaussian smoothing window sizes. Increasing the smoothing scale progressively suppresses transient peak features, including biologically relevant peaks at approximately 2 and 36 hours.

#### Downstream Analysis and Output

SPIFEE provides automated downstream analysis of extracted features, along with a suite of visualization tools designed to support interpretation across multiple levels. Outputs are generated at three primary levels of aggregation: per-event (individual peaks), per-trace, and per-condition summaries. This structure enables both fine inspection of individual signal features and high-level comparisons across experimental conditions.

Core outputs are generated automatically and include quality control diagnostics, peak-level feature tables, condition-level summary statistics, averaged temporal traces, clustering analyses, and statistical comparisons. These outputs are designed to provide a comprehensive overview of signal behavior. In addition, SPIFEE output enables quick, user-driven visualization such as scatterplots and violin plots to facilitate exploratory analysis of relationships between extracted features. The table below summarizes the major output types, their level of aggregation, and their corresponding visualization formats and figure references within this work (Table S2).

| Method | Aggregation Level | Visualization / Format | Reference |
|---|---|---|---|
| ***Core (Automated) Output*** | | | |
| **Quality Control** | Condition | Figures | Figure 1 & Figure 2 |
| **Features of Peaks** | Per Event | Table | Table 3 |
| **Means** | Condition | Table | Not displayed |
| **1st Peaks** | Condition | Table | Not displayed |
| **Average Traces** | Condition | Figure | Figure 8 |
| **Cluster Assignments** | Condition | Figure | Figure 9 A |
| **Trace Clustering** | Condition | Table and Figure | Figure 9 B |
| **Clustering Composition** | Condition or AL | Figure | Figure 9 C |
| **Clustering K Score Evaluation** | Condition | Figure | Figure 9 D |
| **All Condition Clustering** | All Conditions | Table and Figure | Not displayed |
| **Statistics** | Per Feature | Table | Figure 10 |
| **Heatmaps** | Condition | Figure | Figure 11 |
| ***Derived (User-Driven) Output*** | | | |
| **Scatterplots** | Per 2-3 Features | Figure | Figure 12 |

**Table. S2: Overview of automated and user-driven outputs generated by SPIFEE**. Organized by level of aggregation (per-event, per-trace, per-condition, and across conditions). Outputs include tabular summaries, visualizations, and clustering analyses, with corresponding figure references provided where applicable.

#### Average Traces / Basic Feature Extraction

To quantify fluorescence dynamics, SPIFEE extracts features for each detected peak. Core peak features include peak height, location, width, prominence, frequency, and integral. (Table S3). In addition to peak-level features, extracted feature tables also include metadata used for trace indexing and normalization. These include peak number, trace identifier, per-trace summary statistics (e.g., condition average maximum and condition average minimum signal), and derived quantities such as peak-over-basal ratios and total peak counts. While these fields primarily support internal organization and downstream analyses, they are retained in the output for reproducibility and user exploration.

| **Sample Features** | **Trace 1 Peak 1** | **Trace 1 Peak 2** | **Trace 1 Peak 3** |
|---|---|---|---|
| *Height (a.u)* | 400.3 | 277.3 | 284.1 |
| *Location (Hours)* | 12 | 46 | 61 |
| *Width (Hours)* | 2.321 | 1.26 | 2.58 |
| *Prominence (a.u)* | 239.8 | 81.4 | 51.4 |
| *Frequency (Peaks/Trace)* | 0.186 | 0.186 | 0.186 |
| *Integral (a.u)* | 2065.1 | 516.5 | 1629.1 |
| *Peak#* | 1 | 2 | 3 |
| *Trace#* | 1 | 1 | 1 |
| *AvgMax* | 576.2485028 | 576.2485028 | 576.2485028 |
| *AvgMin* | 149.1520225 | 149.1520225 | 149.1520225 |
| *Peak over Basal* | 2.683804913 | 1.859259594 | 1.904841477 |
| *NumPeaks (Trace)* | 3 | 3 | 3 |

**Table S3. Example feature extraction output.** Features are computed for individual detected peaks, including amplitude-based (height, prominence, integral) and temporal (location, width, frequency) characteristics. Additional fields provide trace-level context and bookkeeping information, such as peak index, trace identifier, and baseline averages.

SPIFEE also computes and plots simple condition-level average trace within each condition. (Fig. S7).

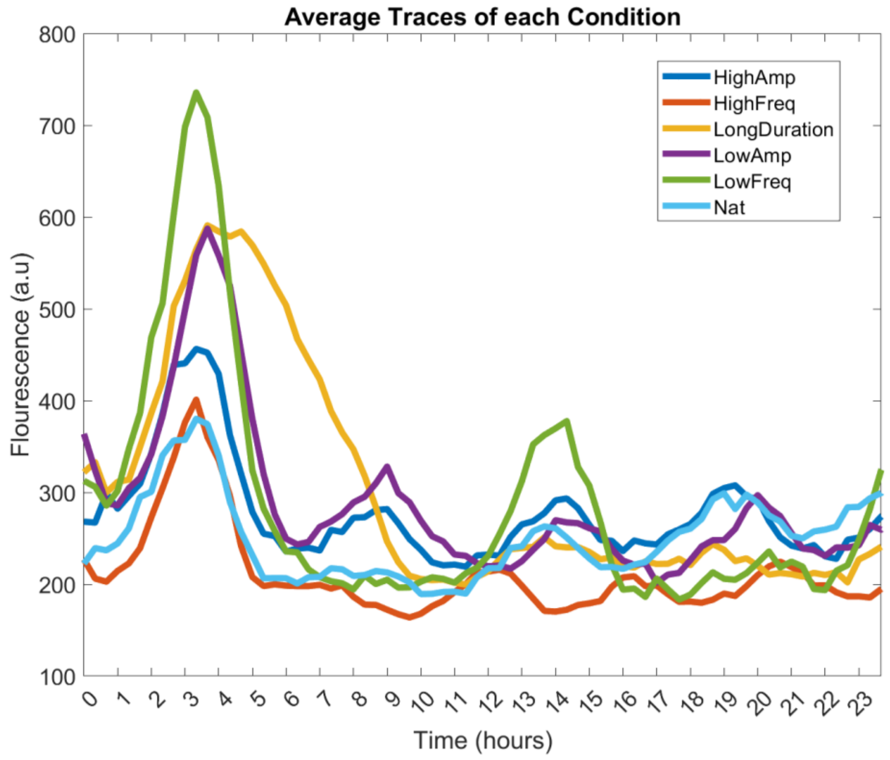

**Fig. S7: Average fluorescence traces for each experimental condition**. These traces correspond to the Nutlin-3–modulated p53 data, illustrating the aggregate dynamics of each condition.

### Clustering

SPIFEE includes a suite of clustering and visualization tools to support identification of subpopulations within fluorescence datasets (Fig. S8). Traces are grouped using k-means clustering with a user-defined distance metric. To guide selection of the number of clusters ($k$), SPIFEE computes four standard clustering evaluation metrics: the Calinski–Harabasz index, Davies–Bouldin index, gap statistic, and silhouette score.

Clustering results are visualized through multiple outputs. Individual trace assignments are displayed to illustrate grouping structure, while cluster-average traces summarize the dynamics of each cluster. In addition, cluster composition plots show the relative proportion of traces assigned to each group, providing insight into the prevalence of distinct response patterns.

Another option available to users is the "Cluster All" option that combines all of the input files and runs clustering on the aggregated data, showing data trends across conditions.

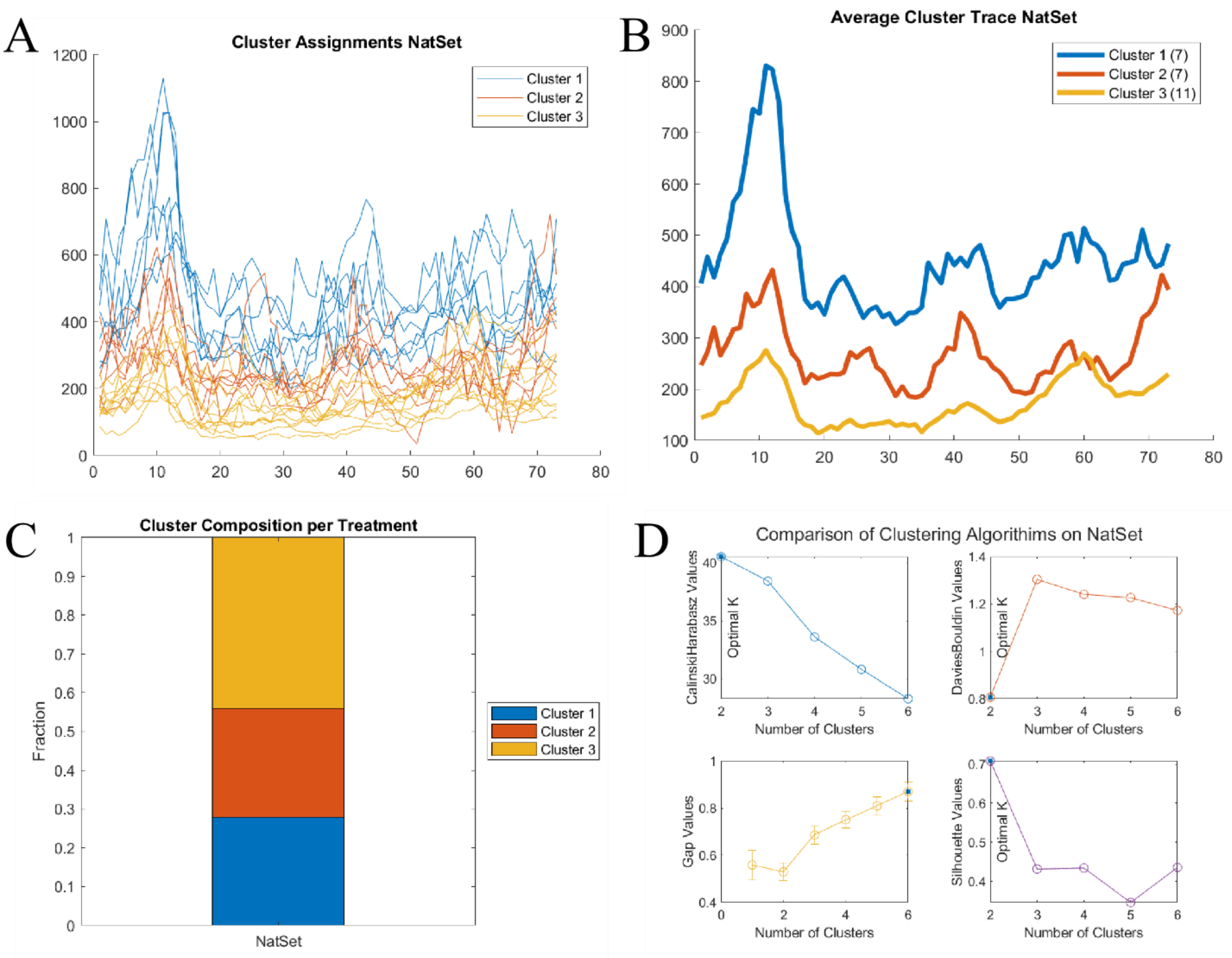

**Fig. S8: Clustering Figures**. (A) Example clustering of individual traces within a single experimental condition using k-means clustering, illustrating separation of distinct dynamic behaviors. In this example, one cluster exhibits a strong early peak followed by heterogeneous pulsing, while another shows a more gradual or sustained response profile.
(B) Cluster-average traces summarizing the characteristic temporal dynamics of each group.
(C) Cluster composition, showing the proportion of traces assigned to each cluster.
(D) Evaluation of clustering performance across varying numbers of clusters ($k$) using four metrics: Calinski–Harabasz index, Davies–Bouldin index, gap statistic, and silhouette score.

### Statistics

SPIFEE provides statistical testing to evaluate differences in extracted features across experimental conditions. For each feature, the distribution of values is first assessed for normality using the Kolmogorov–Smirnov test and for homogeneity of variance using Levene's test. Based on these assumptions, SPIFEE automatically selects an appropriate statistical test: one-way ANOVA when both assumptions are satisfied, Welch's ANOVA when variances are unequal, and the Kruskal–Wallis test when normality is violated. In addition to p-values, SPIFEE computes effect sizes for each comparison to quantify the magnitude of observed differences independent of sample size. Multiple hypothesis testing correction is applied where appropriate. SPIFEE includes a significance flag that highlights comparisons meeting user-defined or default significance thresholds. This flag is intended to guide downstream analysis and highlight key differences while avoiding overreliance on p-values alone. (Fig. S9)

| | 1<br>Feature | 2<br>Condition | 3<br>NumDifferences | 4<br>MeanEffect | 5<br>Flag |
|---|---|---|---|---|---|
| 1 | "Frequency" | "P53_LongDuration" | 5 | -0.3421 | "Distinct" |
| 2 | "Width" | "P53_LongDuration" | 5 | 0.4493 | "Distinct" |
| 3 | "Height" | "P53_HighFreq" | 4 | -0.4370 | "Distinct" |
| 4 | "Integral" | "P53_HighFreq" | 4 | -0.4540 | "Distinct" |
| 5 | "Integral" | "P53_LongDuration" | 4 | 0.4131 | "Distinct" |
| 6 | "Prominence" | "P53_LowFreq" | 4 | 0.4777 | "Distinct" |
| 7 | "Height" | "P53_Nat" | 3 | -0.2878 | "Distinct" |
| 8 | "Prominence" | "P53_HighFreq" | 3 | -0.4319 | "Distinct" |
| 9 | "Prominence" | "P53_LowAmp" | 3 | 0.1042 | "Distinct" |
| 10 | "Prominence" | "P53_Nat" | 3 | -0.3999 | "Distinct" |
| 11 | "Height" | "P53_LongDuration" | 2 | 0.3595 | "" |
| 12 | "Height" | "P53_LowAmp" | 2 | 0.2921 | "" |
| 13 | "Height" | "P53_LowFreq" | 2 | 0.4536 | "" |
| 14 | "Integral" | "P53_HighAmp" | 2 | 0.0296 | "" |
| 15 | "Integral" | "P53_LowAmp" | 2 | 0.0499 | "" |

**Fig. S9: Statistically flagged condition features.** Highlights features that differ significantly between conditions, based on statistical testing and corresponding effect sizes.

#### Heatmaps

Finally, SPIFEE generates heatmaps across experimental conditions to visualize trends and variability in feature distributions, showing global patterns and potential outliers (Fig. S10)

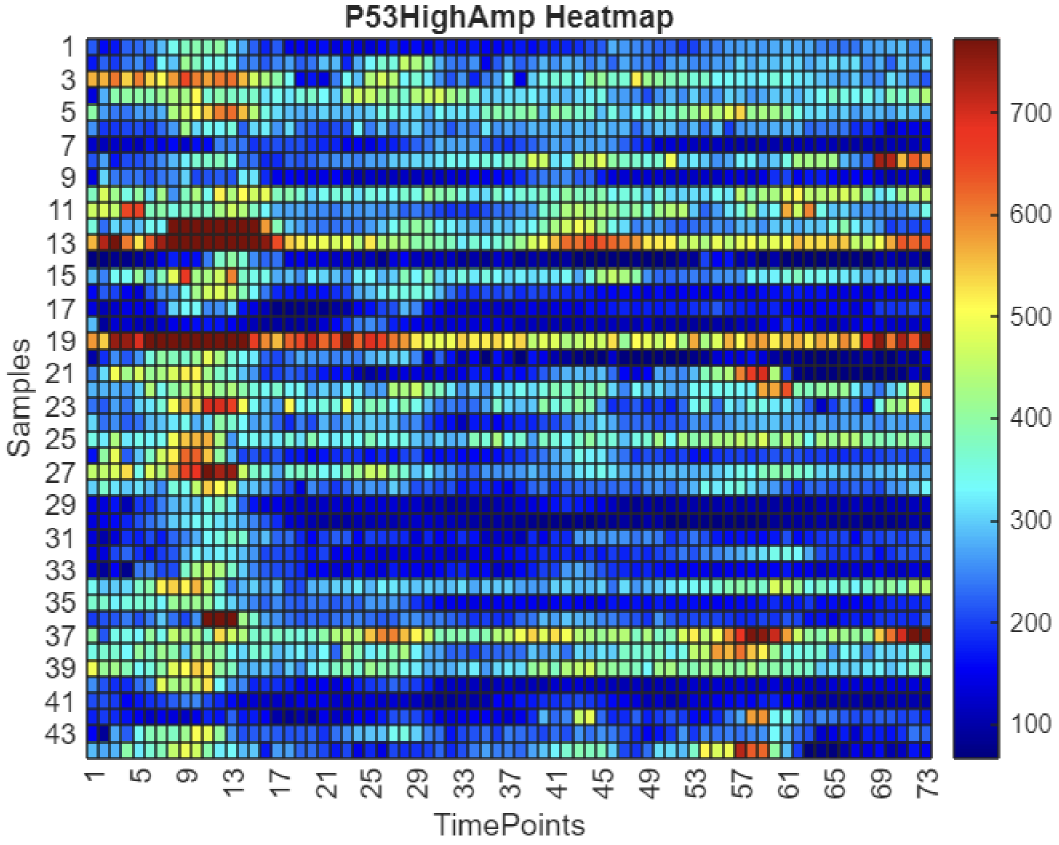

**Fig. S10: Condition-level heatmap.** Fluorescence intensity measurements for each sample are shown across sequential time points. Rows correspond time bins, and columns correspond to samples revealing differences in signal magnitude and temporal progression.

In total, these downstream analysis methods and output data structures provide an accessible and systematic approach for users to explore their data. SPIFEE can be used for new hypothesis generation, to identify hidden subpopulations that may not be apparent through visual inspection, and to assess the impact of different data processing choices.

#### User-Derived Outputs

After utilizing SPIFEE, the output structure is easily parsed to create follow up data visualizations. Here we present one example; a scatter plot of height and width of each peak that was found for 2 conditions from the p53 $H_20_2$ data. (Fig. S11). Inspection of mean values tells you that the average height and width for the 50um treatment is [177.8 (a.u), 1.734 (hours)] and the averages are [292.71 (a.u), 2.058 (hours)] respectively. This doesn't tell the full story, and through this scatterplot, we see how there is quite a lot of overlap in the general peak landscape, and the 80um condition contains far more outliers. This analysis gives insight beyond the population averages and is trival and easy to create using the SPIFEE output structure. Further analysis enabled by SPIFEE include graphs such as violin plots and box charts, among other options.

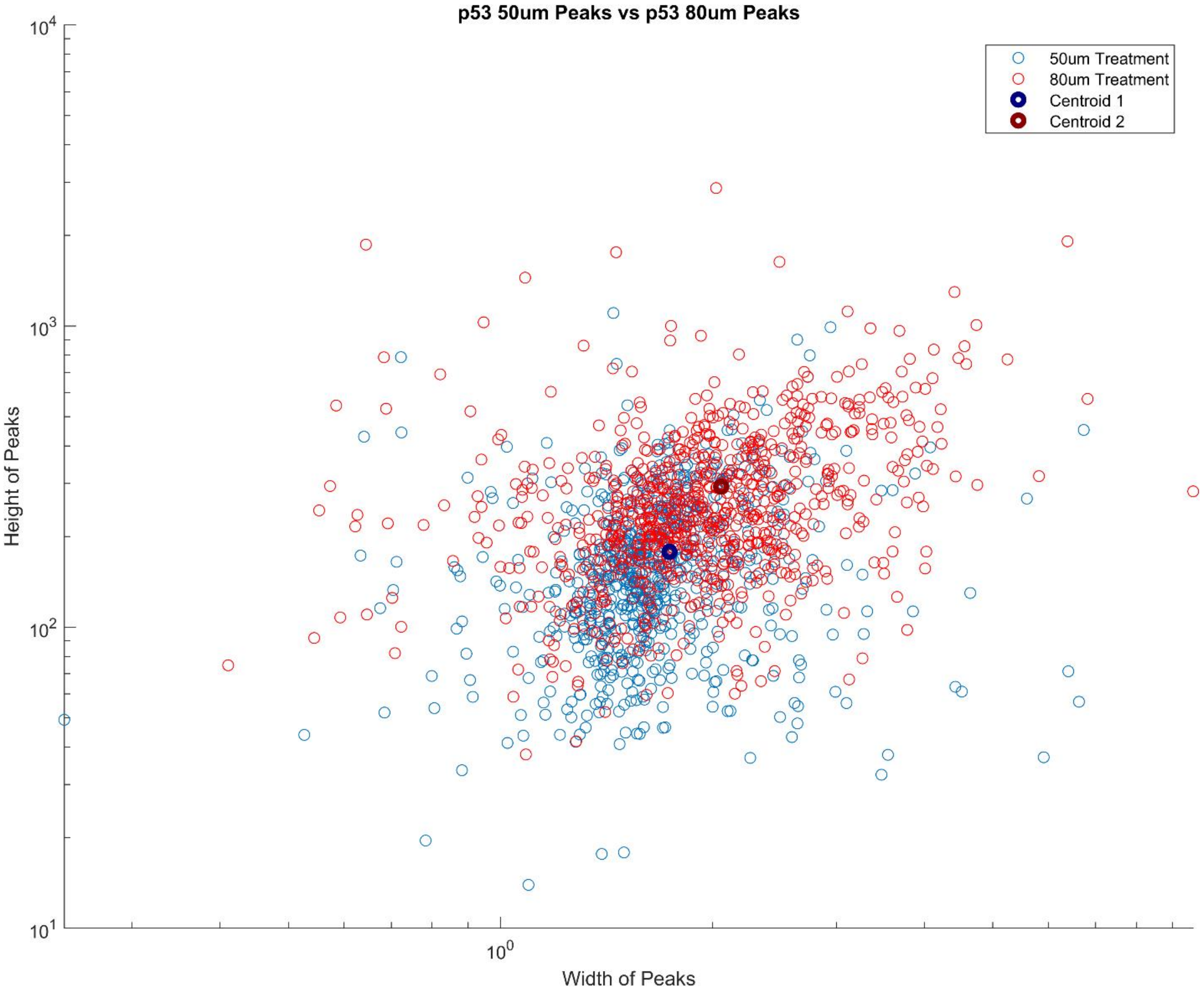

**Fig. S11: Example downstream plot of SPIFEE features.** Widths and heights of peaks are plotted to visualize the centroid output from the clustering, and to visualize the relative feature landscape between the two $H_2O_2$ experiments. Centroids are also calculated and visualized to show global trend.

### Limitations of the Pipeline

While more sophisticated analytical approaches such as wavelet analysis or advanced Fourier transformations exist(2), the goal of this pipeline is not to reinvent existing methodologies. Our aim is to provide an accessible and standardized framework that can be readily applied across experimental conditions and setups, serving as a robust first pass analysis for researchers.

The primary limitations of the pipeline stem from inherent challenges in the data itself:

1. **Filtering and feature detection**: While our filtering and peak detection methods are supported by extensive testing, they are necessarily heuristic and cannot be entirely objective. Users should exercise judgment when interpreting borderline events.
2. **Incomplete or missing data**: SPIFEE requires complete traces for optimal performance. NaN values can be imputed using MATLAB's fillmissing() with the "endvalues" method, but such imputations require careful consideration. By default, traces containing NaNs are excluded.
3. **Temporal scale sensitivity:** The pipeline performs best on oscillatory signals with dynamics on the timescale of hours. Slowly varying signals, traces with very shallow peaks, or very long time-series (beyond ~7 days) are not optimal.
4. **Data resolution / noise**: SPIFEE cannot reliably detect transient pulses in traces with insufficient temporal resolution or in traces with an extreme amount of noise.
